# Green Lacewing *Chrysoperla rufilabris* (Neuroptera: Chrysopidae) is a potential biological agent for controlling crapemyrtle bark scale (Hemiptera: Eriococcidae)

**DOI:** 10.1101/2022.04.18.488594

**Authors:** Bin Wu, Runshi Xie, Mengmeng Gu, Hongmin Qin

## Abstract

Crapemyrtle bark scale (CMBS; *Acanthococcus lagerstroemiae*), an invasive sap-sucking hemipteran, has spread across 16 U.S. states. Infestation of CMBS negatively impacts the flowering of crapemyrtles and reduces the aesthetic quality of ornamental plants. The widespread use of soil-applied neonicotinoid insecticides to repress the CMBS infestation could threaten important beneficial insects; therefore, using natural enemies to control CMBS is greatly needed. This study evaluated larval green lacewing (*Chrysoperla rufilabris*) as a biocontrol agent of CMBS. Predatory behavior of the larval *C. rufilabris* upon CMBS was documented under a stereomicroscope using infested crapemyrtle samples collected from different locations in College Station. Predation potential of *C. rufilabris* upon CMBS eggs and foraging performance using Y-maze assay were both investigated in laboratory conditions. Results confirmed that larval *C. rufilabris* preyed on CMBS nymphs, eggs, and adult females. The evaluation of predation potential results showed that the number of CMBS eggs consumed in 24 hours by 3^rd^ instar *C. rufilabris* (176.4 ±6.9) was significantly higher than by 2^nd^ instar (151.5±6.6) and by 1^st^ instar (11.8±1.3). The foraging performance results showed that larval *C. rufilabris* could target CMBS under dark, indicating that some cues associated with olfactory response were likely involved when preying on CMBS. This study is the first report that validated *C. rufilabris* as a natural predator of CMBS and its potential as a biological agent to control CMBS. Future investigation about the olfactory response of larval *C. rufilabris* to CMBS would benefit the development of environmental-friendly strategies to control CMBS spread.

## INTRODUCTION

As an invasive sap-sucking hemipteran initially found on crapemyrtle (*Lagerstroemia* sp.) in Richardson, TX, crapemyrtle bark scale (CMBS; *Acanthococcus lagerstroemiae*) has spread across 16 U.S. states [1–5]. Sooty mold accumulation resulting from feeding and honeydew secretion of CMBS leads to reductions in growth and blooming vigor of host plants [6] and even branch die-back [7], which negatively impacts the crapemyrtle industry in the U.S. Besides on its primary host, increasing observations of CMBS infestation were reported on other economically important plants [8–11] and native species [7, 9, 12], indicated that the CMBS is a polyphagous invasive insect and poses a great risk to the Green Industry [13, 14] and ecosystems [15, 16] in the U.S.

The effectiveness of bark spraying insecticides is limited in controlling CMBS due to (1) the ability of CMBS to shelter under plant crevices and suck phloem-sap of hosts, (2) the protective wax coverings secreted by adult females and late-instar males of CMBS, and (3) its high fecundity [2, 17, 18]. Neonicotinoids systemically applied through soil drench are effective in suppressing CMBS [6]. However, crapemyrtle is an important pollen source for native and non-native bees in the U.S. [19–21] from late spring to early fall [22, 23], especially when other resources are scarce. The neonicotinoids’ negative impacts on pollinators that rely on crapemyrtle pollen are of great concern [18, 24, 25]. Hence, environmentally friendly and efficacious non-chemical alternatives for CMBS management, including plant resistance breeding and biocontrol agents, are needed [9]. To date, cactus lady beetle (*Chilocorus cacti*) is the only biocontrol agent confirmed in laboratory conditions as a predator of CMBS [7, 26]. Given the relatively broad host range of CMBS and the limited pest management strategies, it is imperative to evaluate other potential biocontrol agents against this invasive pest in the U.S.

*Chrysoperla rufilabris* is a common green lacewing in many horticultural and agricultural cropping systems throughout much of the United States [27–29]. The larvae of *C. rufilabris* are generalist predators of various soft-bodied arthropods with a relatively high prey searching and consumption capacity [29–32]. In practice, the *C. rufilabris* larvae have been applied to control *Aphididae* [33, 34] and *Heliothis* spp. [35] in important crops [31, 36, 37]. The foraging efficiency of *Chrysoperla carnea* or lady beetles upon preys were determined by various cues [38–44]. However, little information is available regarding the predation behavior of *C. rufilabris* on CMBS. To validate whether *C. rufilabris* can be integrated into sustainable CMBS management programs, this study (1) investigated predation activities of the green lacewings upon CMBS in landscape and laboratory conditions, (2) evaluated *in-vitro* predation potential of the green lacewing by different developmental stages; and (3) tested its foraging performance under dark condition.

## MATERIALS AND METHODS

### Collection and Maintenance of Predator and CMBS

Two batches of *Chrysoperla rufilabris* larvae and eggs were purchased from ARBICO Organics™ (Oro Valley, AZ) in June and October 2019. Upon arrival, the individual lacewing larvae and eggs were placed in each VWR®disposable Petri dish (60 mm in diameter) and maintained inside a CONVIRON®-BDR 16 growth chamber (Controlled Environments Ltd., Winnipeg, Manitoba, Canada) at 25±1 °C, 60±5 % RH, and a 16 h L:8 h D photoperiod. The lacewings were provided with eggs and nymphs of CMBS, 20 *μ*L-30 *μ*L droplets of artificial diet [45], and water separately placed on fresh crapemyrtle leaves in the Petri dish (Fig. S1).

Nymphs, adults, and eggs of CMBS were collected from naturally CMBS-infested crapemyrtle plants on Texas A&M University campus in College Station, Texas.

Investigation of Predation Activities upon CMBS in Landscape and Laboratory Conditions The CMBS-infected branches were collected at different Texas A&M University campus locations from April to November 2019 for preliminary landscape research to investigate if *C. rufilabris* are present in plants under CMBS infestation. Nymphs, adults, and eggs of CMBS were randomly distributed in a Petri dish (60 mm in diameter), then one larval *C. rufilabris* was introduced into the same Petri dish to test the predation response of larval *C. rufilabris* to CMBS. The predation behavior was documented under Stemi 2000 stereomicroscope (Carl Zeiss AG, Oberkochen, Germany).

### Predation Potential *in vitro*

Two independent experiments were conducted in June and October 2019, respectively, to evaluate the predation potential of *C. rufilabris* on CMBS. Feeding duration, which refers to the time taken by a *C. rufilabris* to consume the first CMBS egg completely, was recorded under the stereomicroscope from the time when the first egg was captured to the time when the egg was utterly consumed. Number of consumed CMBS, which refers to the number of CMBS eggs in a Petri dish (60 mm in diameter) consumed by an individual larval *C. rufilabris* during a 24-h observation period, was counted with the help of ImageJ (National Institutes of Health, Bethesda, MD). A Petri dish containing approximately 300 fresh CMBS eggs without *C. rufilabris* feeding served as the reference. The Petri dish images before and after feeding were used to accurately determine the number of eggs consumed by *C. rufilabris* in 24 hours.

In June, regardless of its developmental stages, thirteen larval *C. rufilabris* (starved for 24 h beforehand) were individually placed into each Petri dish containing approximately 300 fresh CMBS eggs to test the feeding duration and predation potential. The June experiment was repeated six times using the same thirteen *C. rufilabris*.

In October, twenty larval *C. rufilabris* within the same stage (starved for four hours beforehand) were utilized to investigate the effect of *C. rufilabris* developmental stages (1^st^ instar, 2^nd^ instar, and 3^rd^ instar; Fig. S2) on the feeding duration and predation potential. The October experiment was repeated three times using twenty new larval *C. rufilabris* within the same stage.

### Foraging Performance Test under Dark

To better understand the cues primarily impacting the foraging efficiency of *C. rufilabris*, which provides basic information about the CMBS biocontrol strategies, a primary foraging performance test was conducted using a Y-maze under dark (Fig. 1). The Y-maze consisted of three glass vials, namely loading vial, baited vial, and control vial being joined by a Bel-Art Y-tubing connector (SP Scienceware, Wayne, NJ). Before being fixed with the connector using 1mL pipette tips and Parafilm^®^, ten alive gravid females without ovisacs, ten crawlers and twenty eggs of CMBS were placed in the baited vial, one larval *C. rufilabris* was introduced into the loading vial, and the control vial was vacant.

**Figure 1.**
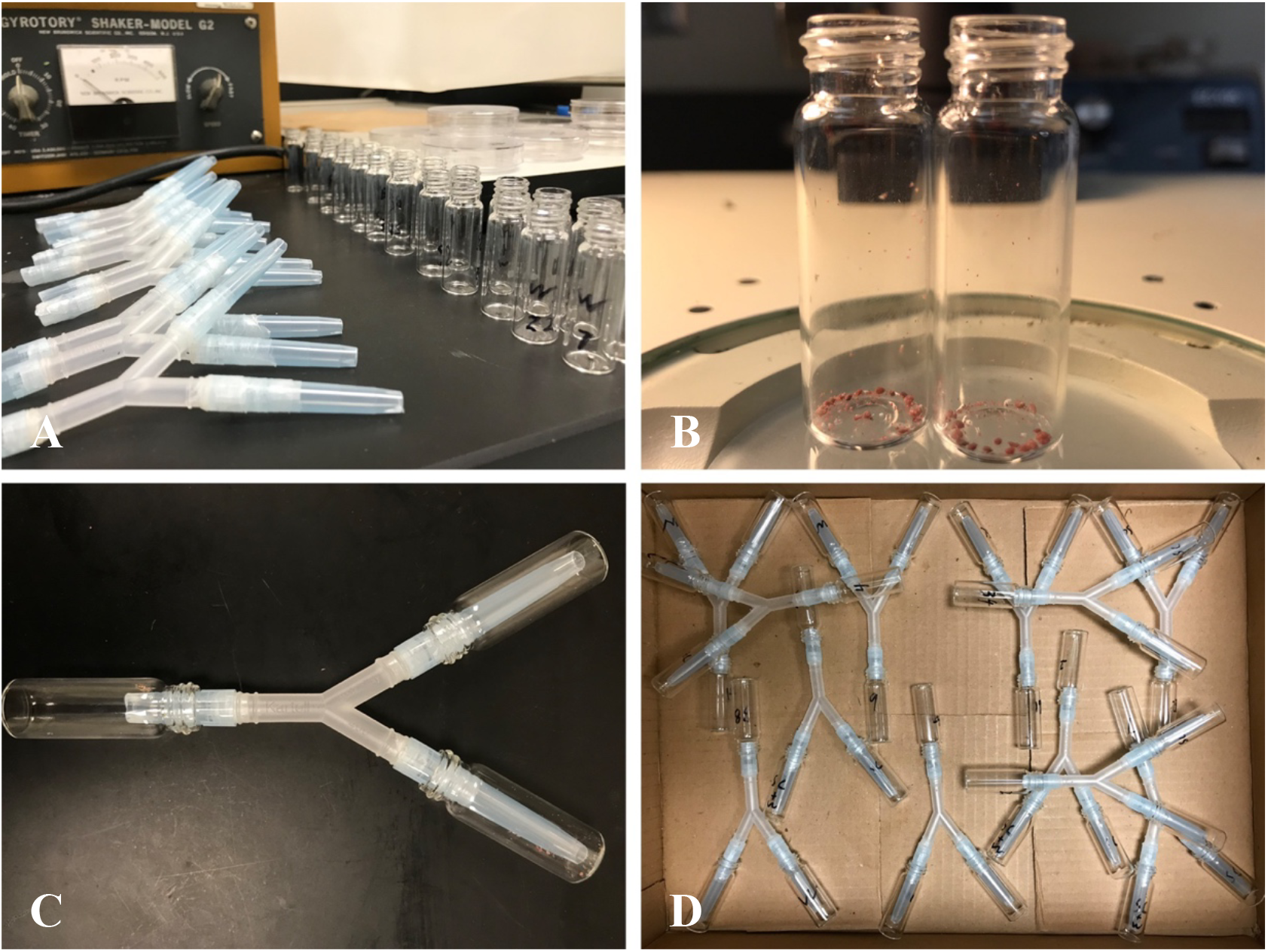
Y-maze assay. A: Each Y-tube setup was assembled by a Y-tubing connector and contained a loading vial, a baited vial, and a control vial; B: Before being fixed to the Y-tubing connector, CMBS females and crawlers were placed into the baited vial; C: Three 1 mL pipette tips wrapped with Parafilm were used to fix the connector to the three vials tightly. Narrow ends of the two pipette tips were cut to connect the baited vial and the control vial, which could avoid the lacewing crawling back once it has made its decision [46]. The wide end of the other pipette tip was cut to connect the loading vial where a larval lacewing was introduced; D: Twelve Y-maze settings were horizontally laid and tested per time in a box and replicated ten times at 25±1°C, 60±5% RH under dark.

Twelve larval *C. rufilabris* were individually placed into each Y-maze setup for the foraging performance test at 25±1°C, 60±5% RH and a 24-h dark photoperiod. After 24 hours, the number of *C. rufilabris* that entered the baited vials (B) and the control vials (C) was counted, respectively. This experiment was repeated ten times using new larval *C. rufilabris*. Foraging performance index (FPI) of *C. rufilabris* targeting CMBS under dark was calculated as:

FPI = (the number of lacewings choosing B - the number of lacewings choosing C) / Total number of lacewings that made a choice Positive response ratio (PRR) of *C. rufilabris* foraging performance was calculated as:

PRR = the number of lacewings choosing B / Total number of lacewings that made a choice

### Data Analysis

For the predation potential experiment in June, datasets of the thirteen *C. rufilabris* biological replicates about the feeding duration and the number of consumed CMBS were averaged (mean ± SE) and compared among the six technical replicates, respectively. For the October experiment, datasets of twenty *C. rufilabris* biological replicates with three technical replicates about the feeding duration and the number of consumed CMBS for each larval stage were analyzed by using one-way analysis of variance (ANOVA) with the JMP®16 (SAS Institute, Cary, NC), respectively. Then, the analysis results regarding the feeding duration and the number of consumed CMBS were separated by *C. rufilabris* developmental stage by using Tukey’s honestly significant difference (HSD; *α* =0.05), respectively, to test if different stages of development impact the *in-vitro* predation potential of larval *C. rufilabris* upon CMBS eggs.

## RESULTS AND DISCUSSION

### Predation Activities of *C. rufilabris* on CMBS Occurred in Landscape and Laboratory Conditions

By examining the samples collected at different campus locations, we observed larval green lacewings feeding on CMBS gravid females [Fig. 2-A&B (30°36’39” N, 96°20’58” W); (30°36’55” N, 96°20’24” W)]. Lacewings’ eggs were deposited on twigs of CMBS-infested crapemyrtles [Fig. 2-C&D, (30°37’03” N, 96°20’08”W); (30°36’30” N, 96°21’02” W)]. These observations encouraged us to evaluate the potential of *C. rufilabris* as a biocontrol agent for sustainable management practices against CMBS. Indeed, in laboratory conditions (Fig. 3), the larval green lacewings not only voraciously consumed CMBS gravid females and eggs but were also able to grab and devour tiny crawling nymphal CMBS. The observations in both landscape and lab conditions confirmed *C. rufilabris* as the natural predator on CMBS.

**Figure 2.**
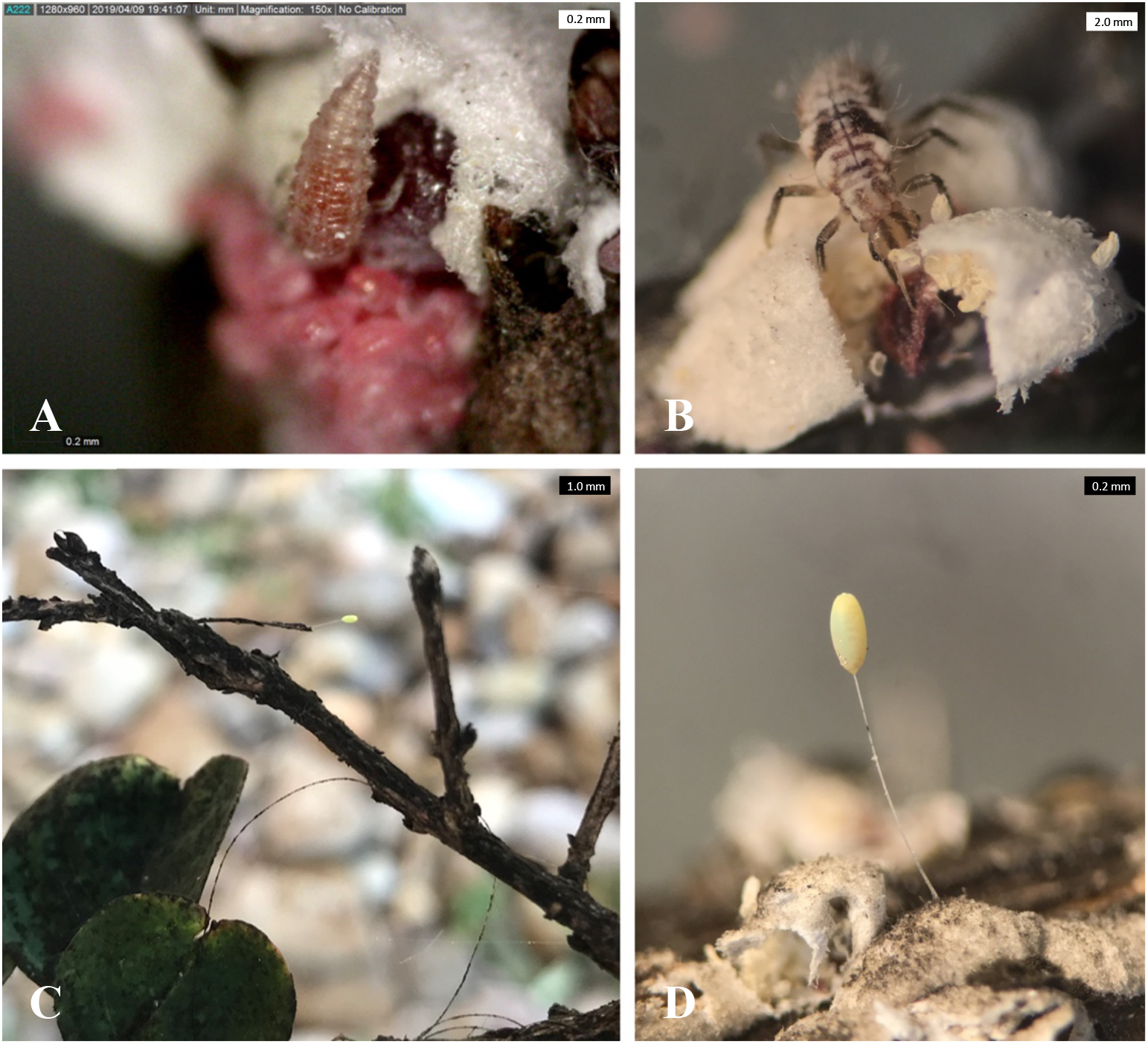
Observations of *Chrysoperla rufilabris* were reported at different locations on Texas A&M campus. Larval *C. rufilabris* were observed preying on CMBS gravid females during the landscape investigations on April 9^th^ (A) (30°36’39” N, 96°20’58” W) and June 28^th^ (B) (30°36’55” N, 96°20’24” W); Lacewing eggs were found in CMBS-infested crapemyrtles on Oct18^th^ (C) (30°37’03” N, 96°20’08” W) and Nov 15^th^, 2019 (D) (30°36’30” N, 96°21’02” W).

**Figure 3.**
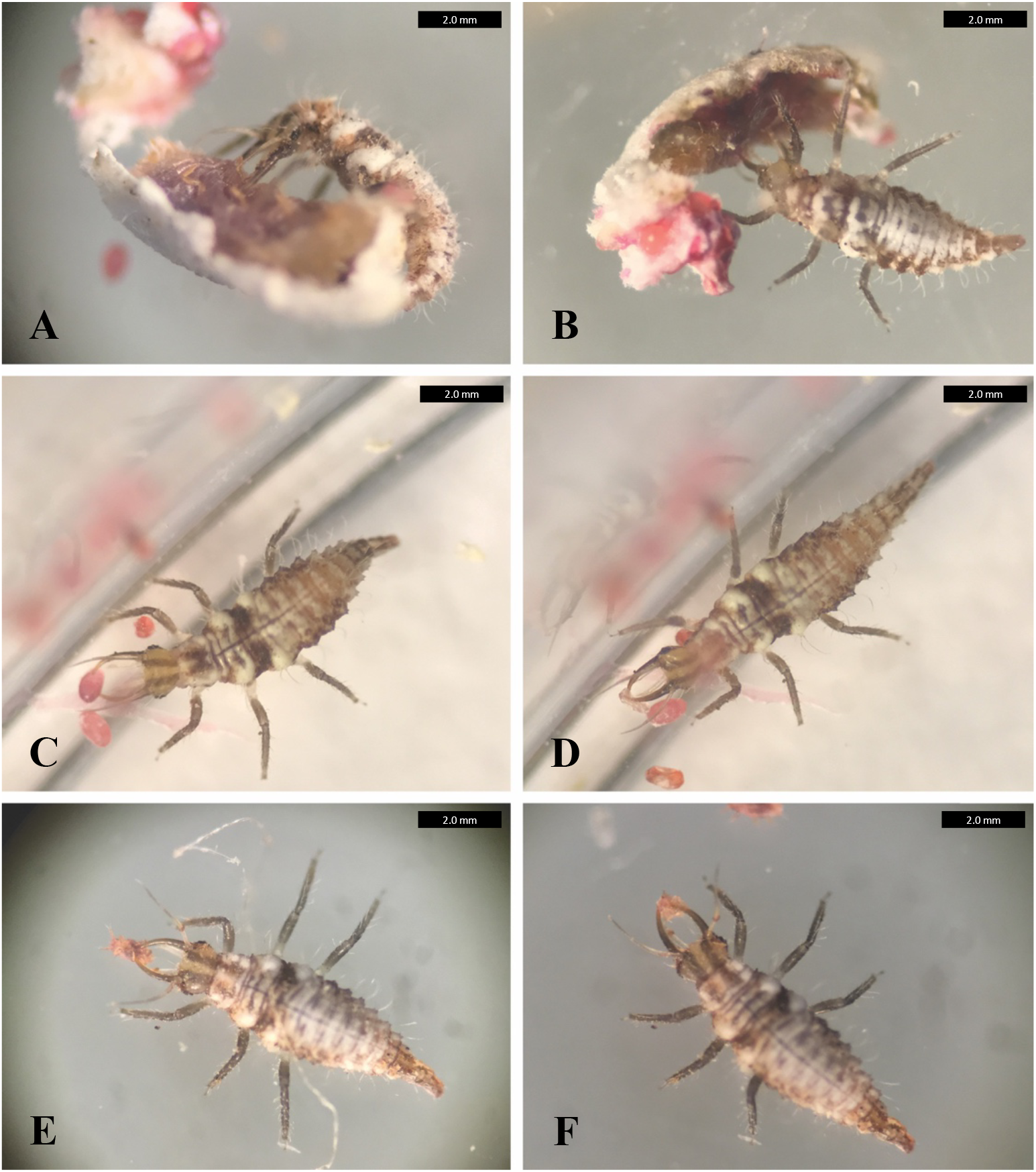
Larval *Chrysoperla rufilabris* individuals were preying on CMBS under laboratory conditions. A larva *of C. rufilabris* targeted a female adult of CMBS (A) and voraciously seized and consumed body fluids of the CMBS using its large, sucking jaws (B) after placing them in the same Petri dish. A green lacewing larva easily grabbed a CMBS egg (C) and consumed the egg in about 1min (D) after placing them in the same Petri dish. Besides, a larva of the green lacewing was able to seize a tiny crawling nymphal CMBS (E) and devour it quickly (F) under the same experiment circumstances.

### Evaluation of the Predation Potential upon CMBS Eggs

In the June test, the results showed that the feeding duration (mean ± SE) ranged from 53.2 ± 2.5 sec to 73.2 ± 2.7 sec and the number of CMBS eggs consumed (mean ± SE) ranged from 154.1 ± 2.7 to 195.5 ± 2.5. The predation potential (or predation capacity) of *C. rufilabris* upon CMBS eggs was similar to that upon fourth-instar aphids of *Aphis gossypii* and *Myzus persicae* [34].

In the October test, the developmental stages significantly affected the feeding duration (*F*_2, 177_ = 101.1332, *p* < 0.0001) and the number of CMBS eggs consumed in 24 h (*F*_2, 177_ = 252.6378, *p* < 0.0001) (Table 1). As predator age increased, the feeding duration of *C. rufilabris* dropped from 141.4 ± 4.8 sec in the 1^st^ instar to 60.3 ± 3.0 sec in the 3^rd^ instar. Meanwhile, the number of CMBS eggs (176.4 ± 6.9) consumed by each lacewing at 3^rd^ stage was significantly higher than the amount (151.5 ± 6.6) consumed by 2^nd^ instar lacewings, followed by 1^st^ instar ones (11.8 ± 1.3). *Chrysoperla rufilabris*, a commercially available biocontrol agent [31], has been validated as the CMBS’s natural predator in this study. The first major peak in CMBS crawler activity occurred in April [47], therefore, to effectively suppress the CMBS population in practice, augmentative releases of the 2^nd^ and 3^rd^ instar *C. rufilabris* during this period should be evaluated further.

**Table 1.** Feeding Duration and Numbers of CMBS Eggs Consumed by *Chrysoperla rufilabris* at Different Developmental Stages

| Predator developmental stage | Feeding duration (sec) <sup>z</sup> | Number of consumed CMBS <sup>z</sup> |
| --- | --- | --- |
| 1<sup>st</sup> instar | 141.4 ± 4.8a | 11.8 ± 1.3c |
| 2<sup>nd</sup> instar | 77.5 ± 4.7b | 151.5 ± 6.6b |
| 3<sup>rd</sup> instar | 60.3 ± 3.0c | 176.4 ± 6.9a |
| Statistical analysis | $F_{2,177} = 101.1332, p < 0.0001$ | $F_{2,177} = 252.6378, p < 0.0001$ |

<sup>z</sup> Means ± SE (N=3, representing a total of 60 tested for each developmental stage), in the same column, followed by different letters are significantly different as determined by Tukey's HSD test ( $\alpha=0.05$ ).

### Foraging Performance Test in Y-mazes

In the 24-h Y-maze assay (Fig. 4-A), the FPI of *C. rufilabris* upon CMBS was 0.56 ± 0.09 (mean ± SE). Among the green lacewings that made a choice, 78.14 ± 4.74% (PRR) larval *C. rufilabris* successfully targeted CMBS in the baited vial under dark (Fig. 4-B). The results indicated that some cues primarily associated with olfactory response were likely involved in the foraging performance. Thus, testing olfactory response to volatiles secreted by preys would help ascertain the attractants or repellents related to lacewing-CMBS interaction, which guides better integrating *C. rufilabris* into the IPM program of CMBS.

**Figure 4.**
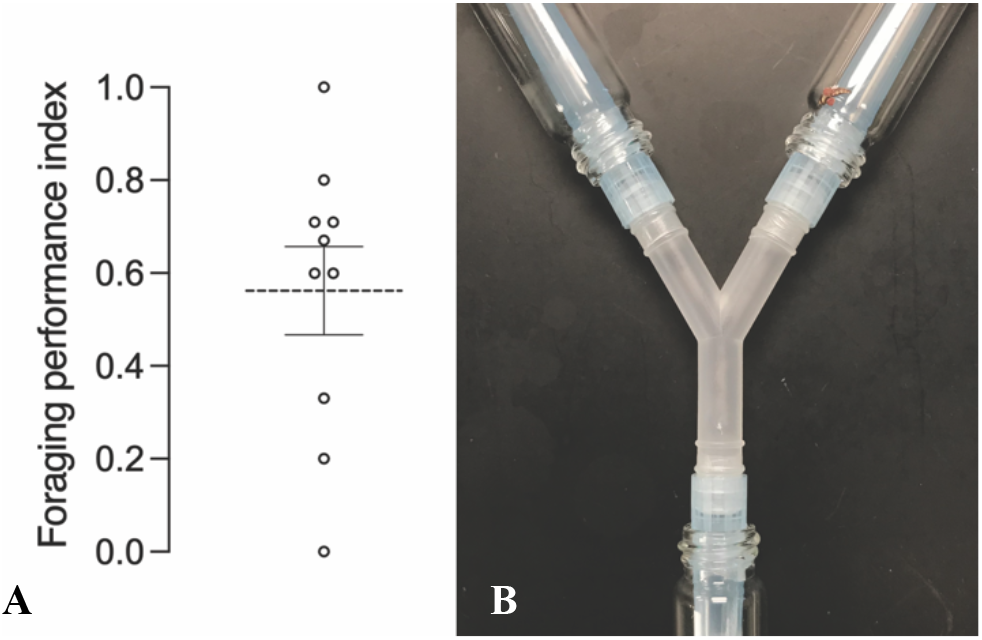
Foraging performance test in 24-h Y-mazes. A: The foraging performance index (FPI) of *C. rufilabris* in the 24-h Y-maze assay was 0.56 ± 0.09 (mean ± SE); B: Among the lacewings that made a choice, 78.14 ± 4.74% larval *C. rufilabris* successfully targeted CMBS under dark.

## CONCLUSIONS

Our study first validated *C. rufilabris* as a natural predator of CMBS and investigated *C. rufilabris* predation potential as a biocontrol agent in laboratory conditions. The results regarding its predation potential upon CMBS eggs suggest using 2^nd^ and 3^rd^ instar *C. rufilabris* could be more efficient in suppressing the CMBS population. The foraging performance results showed that 78.14 ± 4.74% (PRR) larval *C. rufilabris* targeted CMBS under dark, indicating that olfactory response was likely involved in preying on CMBS. Future investigation focusing on the olfactory response of *C. rufilabris* to CMBS would benefit the development of the integrated pest management of CMBS.

## Supporting information

Figure S1 provides a whole picture of rearing the green lacewing Chrysoperla rufilabris. Figure S2 shows the life cycle of C. rufilabris.

## Supplementary Data

Figure S1 provides a whole picture of rearing the green lacewing *Chrysoperla rufilabris*. Figure S2 shows the life cycle of *C. rufilabris*.

## Acknowledgments

This work is supported by TAMU T_3_ 246495-2019, Crop Protection and Pest Management project ‘Integrated pest management strategies for crape myrtle bark scale, a new exotic pest’ (No. 2014-70006-22632) and Specialty Crop Research Initiative project ‘Systematic Strategies to Manage Crapemyrtle Bark Scale, An Emerging Exotic Pest’ (grant no. 2017-51181-26831) from the USDA National Institute of Food and Agriculture. Any opinions, findings, conclusions, or recommendations expressed in this publication are those of the authors and do not necessarily reflect the view of the U.S. Department of Agriculture.

