## Supplementary material for "Green Lacewing *Chrysoperla rufilabris* (Neuroptera: Chrysopidae) is a potential biological agent for controlling crapemyrtle bark scale (Hemiptera: Eriococcidae)": Figure S1 provides a whole picture of rearing the green lacewing Chrysoperla rufilabris. Figure S2 shows the life cycle of C. rufilabris.

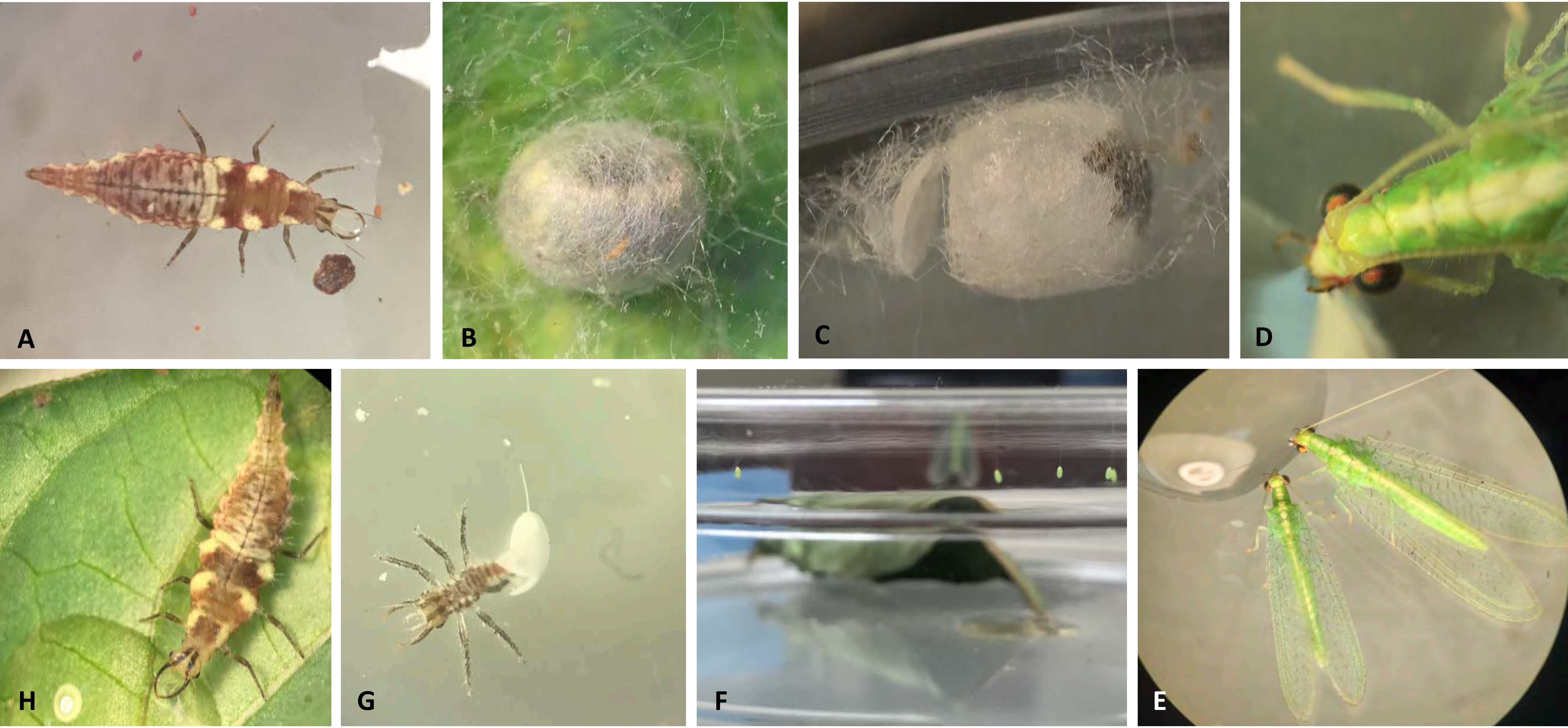


**Supplementary Figure 1**. Rearing *Chrysoperla rufilabris.* Individual larval green lacewings were placed in each Petri dish and fed with eggs and nymphs of CMBS and 20 $\mu$L-30 $\mu$L droplets of artificial diet (Prosser and Douglas 1992) (A). After three larval stages, the third instars secreted silken cocoons to cover themselves to developed as pupae (B). Winged *C. rufilabris* adults emerged from the cocoons (C) and were reared using 20 $\mu$L-30 $\mu$L droplets of artificial diet and water (D). Several *C. rufilabris* adults were transferred into the same Petri dish supplemented with the artificial food for reproduction (E). Green eggs were disposed on the lid of the Petri dish around the food (F). The 1$^{st}$ instar *C. rufilabris* hatched from the eggs (G) and provided with the artificial foods (H).

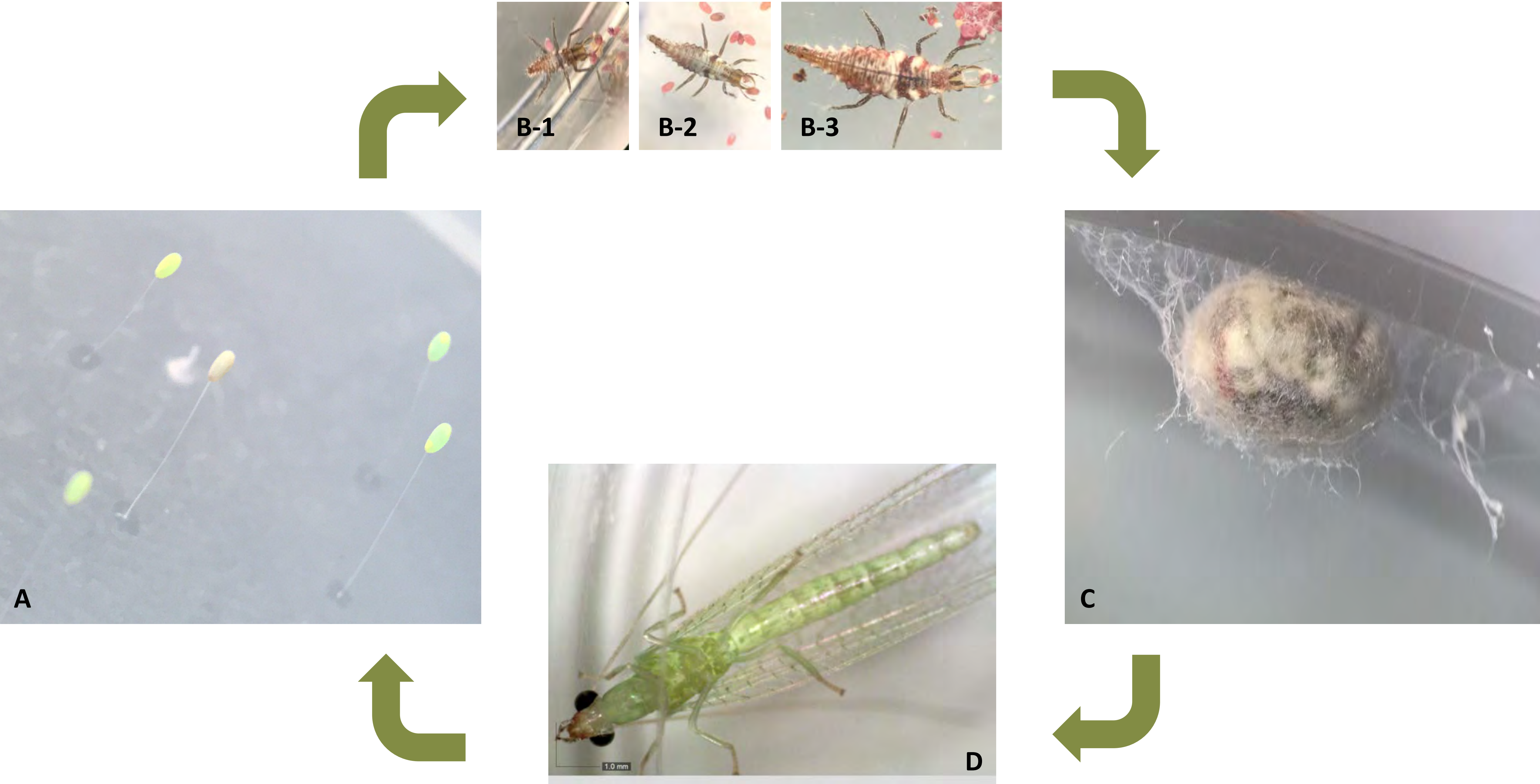


**Supplementary Figure 2**. Life cycle of *Chrysoperla rufilabris.* After hatching from eggs (A) in 5-8 d, the *C. rufilabris* instars developed through 1$^{st}$ (B-1), 2$^{nd}$ (B-2), and 3$^{rd}$ (B-3) larval stages in 14-21 d. Then, the third instars secreted silken cocoons to cover themselves as pupae. Winged *C. rufilabris* adults emerged from the cocoons in 10-15 d.
